# Modulation of immunosuppressant drug treatment to improve SARS-CoV-2 vaccine efficacy in mice

**DOI:** 10.1101/2021.09.28.462156

**Authors:** Amy V. Paschall, Ahmet Ozdilek, Sydney L. Briner, Melinda A. Brindley, Fikri Y. Avci

## Abstract

The COVID-19 pandemic dramatically demonstrated the need for improved vaccination strategies and therapeutic responses to combat infectious diseases. However, the efficacy of vaccines has not yet been demonstrated in combination with commonly used immunosuppressive drug regimens. We sought to determine how common pharmaceutical drugs used in autoimmune disorders can alter immune responses to the SARS-CoV-2 spike protein vaccination.

We treated mice with five immunosuppressant drugs (cyclophosphamide, leflunomide, methotrexate, methylprednisolone, and mycophenolate mofetil), each with various mechanisms of action prior to and following immunization with SARS-CoV-2 spike protein. We assessed the functionality of antibody responses to spike protein and compared immune cell populations in mice that received no treatment with those that received continuous or temporarily suspended immune suppressive therapy.

All tested immunosuppressants significantly reduced the antibody titers in serum and functional antibody response against SARS-CoV-2 spike protein in immunized mice. Temporarily halting selected immunosuppressants (methylprednisolone and methotrexate, but not cyclophosphamide) improved antibody responses significantly. Through proof-of-principle experiments utilizing a mouse model, we demonstrated that immune suppression in autoimmune disorders through pharmaceutical treatments may impair vaccine response to SARS-CoV-2, and temporary suspension of immunosuppressant treatment may be necessary to mount an effective antibody vaccine response. This work provides feasibility for future clinical assessment of the impact of immunosuppressants on vaccine efficacy in humans.

**Significance Statement:** Immunosuppressant regimens are widely used as therapies for a variety of diseases, including autoimmune, inflammatory, and cancer. However, immunosuppressants can impair critical immune responses to vaccination. The impact of standard immunosuppressant use on the critical, developing SARS-CoV-2 vaccination strategies has not been well-described. In this study, we use a mouse model to determine how different immunosuppressant drugs that act through different mechanisms can impair the antibody response to SARS-CoV-2 spike protein, and how modulating these drug regimens may restore antibody levels and function.

## Introduction

In December of 2019, a new severe acute respiratory syndrome beta-coronavirus (SARS-CoV-2) that closely resembled an earlier coronavirus, SARS-CoV-1(1, 2), was identified as the causative agent for the rapidly spreading disease(3), COVID-19. SARS-CoV-2 readily spreads amongst humans(4, 5) and has a mortality rate between two to eight percent(6, 7) in symptomatic individuals. While vaccine efforts against beta-coronaviruses had not been highly prioritized prior to the COVID-19 pandemic, previous research had identified key coronavirus-common antigens for potential vaccine targeting (2). Four common proteins involved in the structure of coronaviruses are the spike protein, membrane protein, nucleocapsid protein, and envelope protein (8). The spike protein includes two subunits: S1 and S2. The S1 subunit contains the receptor-binding domain (RBD)(9), which binds angiotensin converting enzyme 2 (ACE2) on the surface of host cells(5, 10) and is thus necessary for viral entry and propagation. The spike protein and its critical role in the viral infection has been extensively investigated in studies of SARS-CoV-1 and Middle East respiratory syndrome (MERS), and was determined to be highly conserved among human coronaviruses(11, 12). Interfering with spike protein binding and function inhibits virus infectivity, making the spike a highly attractive antigenic target for potential anti-viral treatments, including the currently approved SARS-CoV-2 vaccines and antibody therapies. Antibodies targeting the spike protein can neutralize the virus and prevent its infectivity(2), and are thus a key determinant in determining patient immune protection against SARS-CoV-2.

Most current WHO approved vaccines against SARS-CoV-2 mainly target the spike protein. Studies find the vaccines induce robust antibody response as well as T cell responses that are highly effective at preventing severe disease (13-15). In addition, treatment with cocktails of monoclonal antibodies against spike are effective in treating COVID-19 patients, suggesting antibody levels are an important component of the immune response to SARS-CoV-2. Patients with autoimmune diseases or other diseases that alter the patient immune landscape have been shown to exhibit poorer responses to SARS-CoV-2 vaccination(16-18). Potentially, effective antibody response to SARS-CoV-2 vaccination may be inhibited using immunosuppressants.

Immunosuppressants are commonly used in the treatment of autoimmune diseases, including rheumatoid arthritis and lupus, and are also common requirements following organ transplantation. Immunosuppressants target specific or multiple immune cell populations or functional pathways. Furthermore, cancer chemotherapy drugs may induce immune suppression as a secondary effect Approximately 6 million Americans are estimated to be taking an immune-weakening drug(19). Thus, a large percentage of the population could be expected to generate a weaker immune response following vaccination against COVID-19. Previous research has shown immunosuppressant treatments can significantly inhibit patient responses to vaccinations against multiple pathogens including viral(20, 21) and pneumococcal(22, 23). A number of immunosuppressant drugs have now been shown to inhibit the level of antibody responses and immunogenicity of mRNA vaccines against SARS-CoV-2 in human patients (24). Of note, B-cell depleting therapies showed particularly significant reductions in antibody titers, though other immunocompromising agents showed decreased antibody response as well. These studies indicate the need for optimization of vaccination strategies in immunocompromised patients, more specifically how to induce sufficient antibody titers and prime immune cells against SARS-CoV-2 without quitting immunosuppressant treatments.

In this study, we investigated the effects of five widely used immunosuppressant drugs that target different pathways of the immune responses, and if these effects could be modulated following changes to immunosuppressant regimens in mouse models (Table 1). Specifically, we investigated: cyclophosphamide (CYC, a potent drug known to deplete immune cells, which has been shown to impair vaccine responses(25)); leflunomide (LEF, a disease-modifying antirheumatic drug, or DMARD, that interferes with immune cell replication and has been shown to inhibit IgE antibody response (26, 27)); methotrexate (MTX, a potent drug used in multiple disease models which inhibits the activities of multiple enzymes critical for immune cell function and has been shown to interfere in multiple vaccination models (22-24, 28, 29)); methylprednisolone (MP, a commonly used corticosteroid that may alter opsonophagocytic killing of pathogens by immune cells(23)); and mycophenolate mofetil (MM, an inhibitor of purine biosynthesis shown to inhibit antibody responses (30)). We determined that all drugs significantly reduced antibody response to SARS-CoV-2 spike protein vaccination in mice when administered in doses and routes of administration based on established mouse models (22-30) (Table 1). However, temporary suspension of drugs at vaccination timepoints may improve vaccine response to the spike protein. These findings may guide clinical studies to demonstrate that immunosuppressant administration can be modulated to improve immune responses generated through vaccinations against COVID-19 in humans.

**Table 1.** Immunosuppressants administered to mice.

| **Treatment** | **Trade Name(s)** | **Mechanism of Action** | **Uses** | **Refs** |
| --- | --- | --- | --- | --- |
| Methylprednisolone (MP) | Medrol,<br>DepoMedrol,<br>SoluMedrol | Glucocorticoid steroid | Autoimmune | [18] |
| Methotrexate (MTX) | Trexall, Rasuvo,<br>Otrexup | Antimetabolite<br>(inhibits DHFR) | Autoimmune;<br>chemotherapy | [17-19],<br>[23-24] |
| Cyclophosphamide (CYC) | Cytoxan, Neosar | Metabolite crosslinks with DNA strands → cell apoptosis | Autoimmune;<br>chemotherapy | [20] |
| Leflunomide (LEF) | Arava,<br>Lefumide,<br>Arabloc | DMARD | Autoimmune | [21-22] |
| Mycophenolate mofetil (MM) | CellCept,<br>Myfortic | Inhibits IMPDH to prevent purine synthesis | Autoimmune | [25] |

Table 1.
| Treatment | Trade Name(s) | Mechanism of Action | Uses | Refs |
| --- | --- | --- | --- | --- |
| Methylprednisolone (MP) | Medrol, DepoMedrol, SoluMedrol | Glucocorticoid steroid | Autoimmune | [18] |
| Methotrexate (MTX) | Trexall, Rasuvo, Otrexup | Antimetabolite (inhibits DHFR) | Autoimmune; chemotherapy | [17-19], [23-24] |
| Cyclophosphamide (CYC) | Cytoxan, Neosar | Metabolite crosslinks with DNA strands → cell apoptosis | Autoimmune; chemotherapy | [20] |
| Leflunomide (LEF) | Arava, Lefumide, Arabloc | DMARD | Autoimmune | [21-22] |
| Mycophenolate mofetil (MM) | CellCept, Myfortic | Inhibits IMPDH to prevent purine synthesis | Autoimmune | [25] |

## Results

### Immune suppression impairs antibody response to SARS-CoV-2 spike protein in primary and boost immunizations

We first sought to determine whether commonly used immune suppressants may dampen the antibody responses to SARS-CoV-2 spike protein, possibly reducing COVID-19 vaccine efficacy. We therefore started mice on immune suppression regimens of CYC, LEF, MM, MP, and MTX, as well as no drug control groups that received phosphate buffered saline (PBS). Each of these immune suppressants has been used in treatment of autoimmune disorders as well as in research of immune responses (Table 1). To emulate clinical spike protein-based vaccination strategies inducing adaptive humoral immunity, we immunized mouse groups with 0.5 μg/mouse of SARS-CoV-2 recombinantly expressed spike protein in alum, or an alum only control, 7 days after starting treatment regimen (Day 0), and then immunized again 14 days after primary immunization (Fig 1A). We first analyzed serum IgG titers at 14 days post-primary and days 17, 21, 28, and 35, which followed booster immunization (Fig 1B). All drug treatment groups showed significant decreases in absorbances compared to those of the No Drug control mouse samples. We analyzed individual serum IgG (Fig 1C) and IgM (Fig 1D) titers following booster immunization (Day 21) as well as the endpoint of the immunization experiment (Day 35) and observed significant differences in IgG titers in all immunosuppressant-treated mouse groups. Similar trends were observed in IgM titers. These results indicate that immunosuppressant regimens can impair the antibody response to SARS-CoV-2 spike protein vaccination. To determine if this inhibition of IgG and IgM titers against spike protein would translate into functional deficiencies in anti-spike response specifically, we utilized a recombinant vesicular stomatitis virus (rVSVΔG) lacking the VSV glycoprotein and encoding the SARS-CoV-2 spike to monitor virus neutralization (Fig 2). When comparing 50% neutralization activity 28 days following primary vaccination, we observed the drug treated sera was not as potent as the No Drug control group (No Drug) except for LEF which had no significant differences in activity compared to control. CYC treatment group did not contain any detectable neutralizing activity at any timepoint. For the endpoint (Day 35), on average, MTX required 8 times more sera than the control group to neutralize 50% of the virus and MP treatment required 9 times as much serum (Fig 2B). We observed functional inhibition of antibody-mediated clearance of spike protein-expressing cells in nearly all of the immunosuppressant-treated groups as compared to control, indicating that continuous immunosuppressant regimens can effectively reduce vaccination response against SARS-CoV-2.

**Figure 1.**
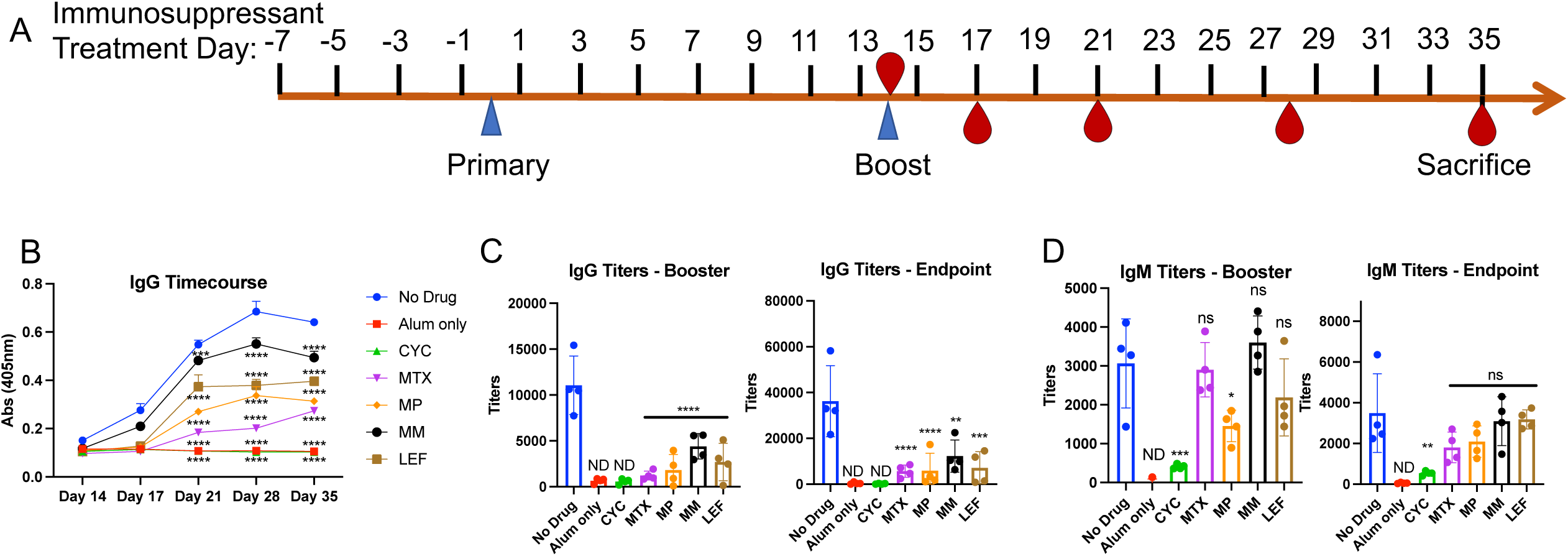
Changes in antibody levels to SARS-CoV-2 spike protein following immunosuppressant administration. **A**. Schematic of intraperitoneal SARS-CoV-2 spike protein immunization and immunosuppressant drug administration. Recombinant spike protein was injected at 0.5μg/50μl alum per mouse. Alum only mice received 50 μl of alum with no spike protein. Immunosuppressants were administered intraperitoneally at the indicated timepoints according to the concentrations in Supplementary Table 1. **B**. Serum was obtained from immunosuppressant-treated mice on days 14 (prior to booster immunization), 17, 21, 28, and 35 days (following booster immunization). ELISA was performed to determine serum IgG titers against spike protein. We did not detect IgG/IgM response to spike protein in the non-immunized mouse sera (alum only). Input was individual OD (405nm) values of sera at 1/3200 dilution and significance was determined for all time points following booster immunization. **C**. IgG and **D**. IgM titers after booster immunizations (day 21) and at endpoint (day 35). Averages of OD values at 405nm absorbance of technical replicates for individual mouse serum sample were used in regression curves to obtain titers (reciprocal serum dilutions) at absorbance of 0.5 at 405nm. Significance was determined as compared to No Drug control group. Alum only and CYC groups were non-detectable (ND).

**Figure 2.**
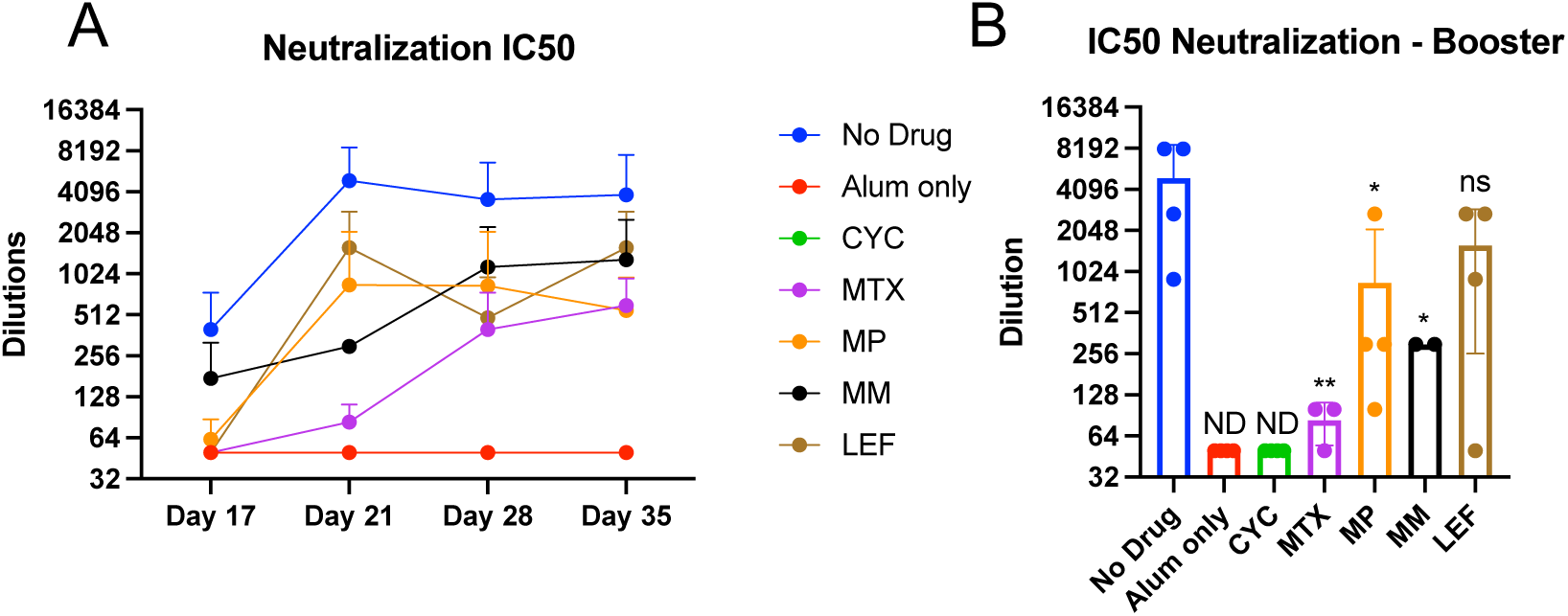
Changes in antibody neutralization efficacy in a VSV-SARS-CoV-2 spike protein system. **A**. Individual serum samples (n=4) were collected from immunosuppressant-treated mice at 17, 21, 28, and 35 days following immunization with spike protein. Sera was serially diluted and mixed with rVSVΔG/SARS-CoV-2-S-NLucP particles. Infectivity of sera incubated virus was assessed on three replicate wells of Vero cells 24 hour following infection by quantifying the levels of NLuc produced. The serum dilutions that limited 50% of the NLuc signal produced by virus lacking sera were plotted. B. Neutralization using booster serum (Day 21) from immunosuppressant mice was compared to serum from No Drug control mice. Noted dilutions are plotted on a Log2 scale.

### Temporary suspension of immunosuppressant regimens is sufficient to recover antibody generation and function against SARS-CoV-2 spike protein

We next sought to determine if this inhibition of an effective response to spike protein immunization in an immunosuppressant regimen could be recovered through temporary suspension of the immunosuppressant administration. To test this, we treated BALB/c mice regularly with three of the immunosuppressant drugs used in this study (CYC, MTX, and MP), halting treatments at the time of immunization or at the timepoints surrounding immunization as indicated in Fig 3A. The three drugs selected for this experiment were based on the consistently lower antibody responses upon their administration compared to MM and LEF as indicated in Fig1B-D and Fig2. This reduced number of immunosuppressant drugs was chosen to also minimize potential experimental variations in further experiments (i.e. plate-to-plate variability). We immunized with spike protein 8 days after the immunosuppressant regimens started and boosted 14 days after primary immunization. We obtained serum at day 14, immediately before the booster immunization, and days 17, 21, 28, and 35, which followed the booster immunization, and performed ELISAs to compare IgG titers in mouse groups that received continuous immunosuppressant treatments through immunization and mouse groups that were temporarily untreated at or around the time of immunization. We observed that all mouse groups that received CYC treatments retained negligible levels of IgG and IgM titers against spike protein, despite interruption of the treatment regimen for both one day and three days of drug suspension (Fig 3B). MP, however, exhibited a partial restoration of antibody response as indicated in ELISA, with the highest level of antibody titers observed in mice that did not receive MP two days before, the day of, and two days after each vaccination timepoint (Fig 3C). We also observed significant increases in booster titer levels when MTX treatment was halted for 1 or 3 timepoints, compared to continuous MTX treatment. This data indicates that antibody response to SARS-CoV-2 spike protein can be improved through modulation of the immunosuppressant regimen, depending on the drug in use.

**Figure 3.**
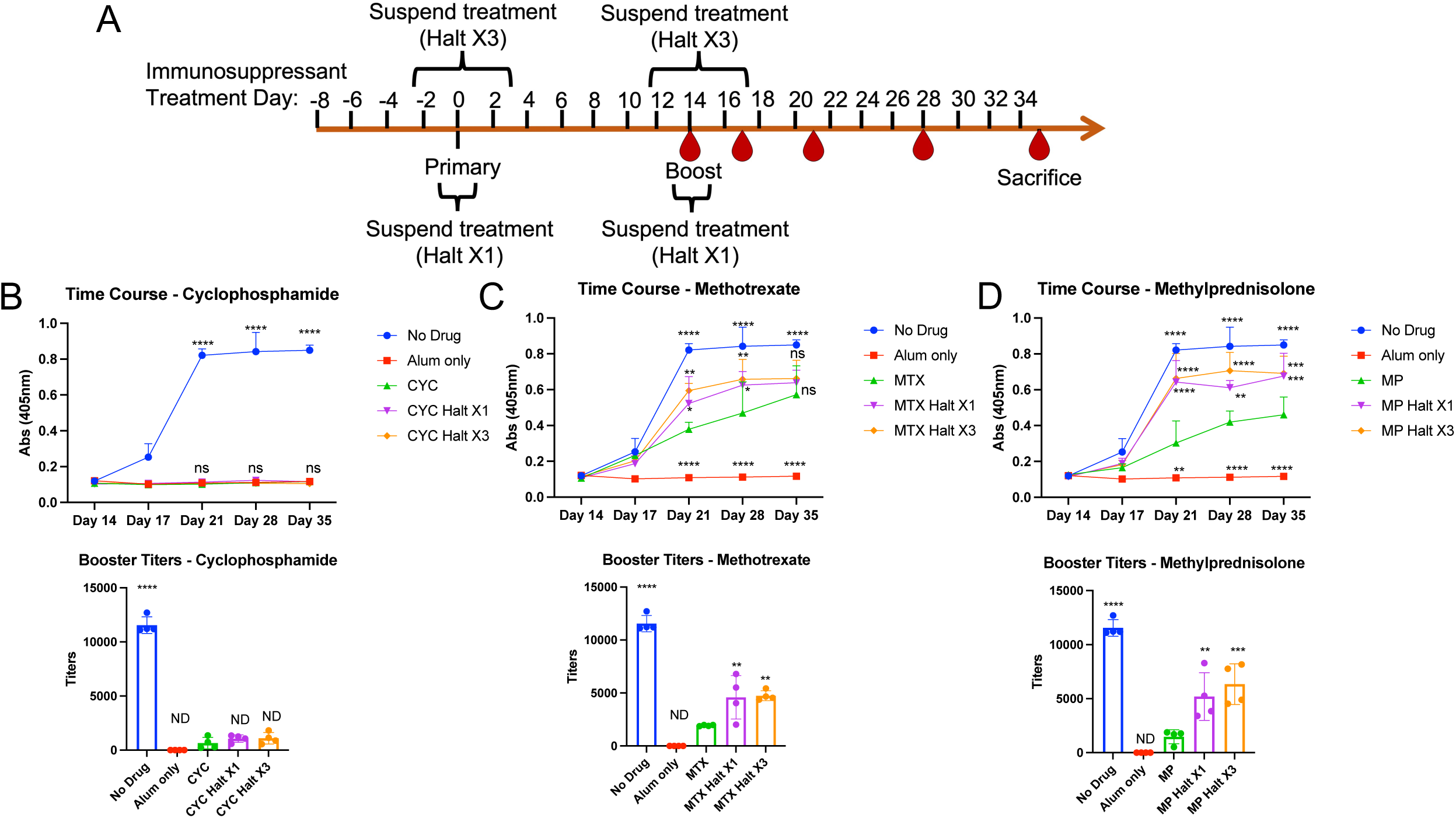
Changes in antibody titers to SARS-CoV-2 spike protein following continuous or temporarily-suspended immunosuppressant administration. **A**. Schematic of intraperitoneal SARS-CoV-2 spike protein immunization and immunosuppressant administration. Spike protein was injected at 0.5μg/50μl alum per mouse. Immunosuppressants were administered intraperitoneally at the indicated timepoints according to the concentrations in Supplementary Table 2. Within each immunosuppressant group, 4 mice did not receive the respective immunosuppressant on days 0 and 14, and 4 mice did not receive immunosuppressants on days - 2, 0, 2, 12, 14, and 16. **B-D**. Serum was obtained from immunosuppressant-treated mice at days 14, 17, 21, 28 and 35. ELISA was performed to determine serum IgG against spike protein. Multiple serial serum dilutions were performed, with each dilution’s set of samples (including all timepoints) on one 384-well plate. Input was individual OD values at 405nm absorbance. Shown are ELISA data using 1:3200 serum dilutions from 4 mice/group (top panels) and calculated booster titers (Day 21) based on regression curves from serial dilutions (bottom panels). Statistical values were determined as compared to continuous drug treatment (A. CYC; B. MTX; C. MP).

Furthermore, this translated to functional differences as demonstrated by the level of sera required to neutralize rVSVΔG-SARS-CoV-2-S infectivity (Fig 4). While halting CYC treatments at or around the time of vaccination did not increase neutralization of the viral particles above the level observed in the continuous CYC treatments (not detectable, Fig 4A), temporary suspension of MP showed higher neutralization compared to the respective continuous treatments (Fig 4C). This indicates that a functional antibody response against the spike protein can be improved through modulation of certain immunosuppressant drugs. While a trend in increased neutralization in the MTX Halt groups (greater than 5X dilution in average comparing Halt X3 to continuous MTX treatment) was observed, this was not considered significant in statistical analysis (Fig 4B). In an additional statistical evaluation, effect size calculation was performed with the Halt X1 and X3 groups compared to continuous drug treatments (Supplementary Table 6). Halting both MTX and MP for either X1 or X3 timepoints showed very large effect size of the standardized mean differences using Cohen’s d calculation compared to the respective continuous drug regimens. These data indicate that a functional antibody response against the spike protein can be improved through modulation of certain immunosuppressant drugs.

**Figure 4.**
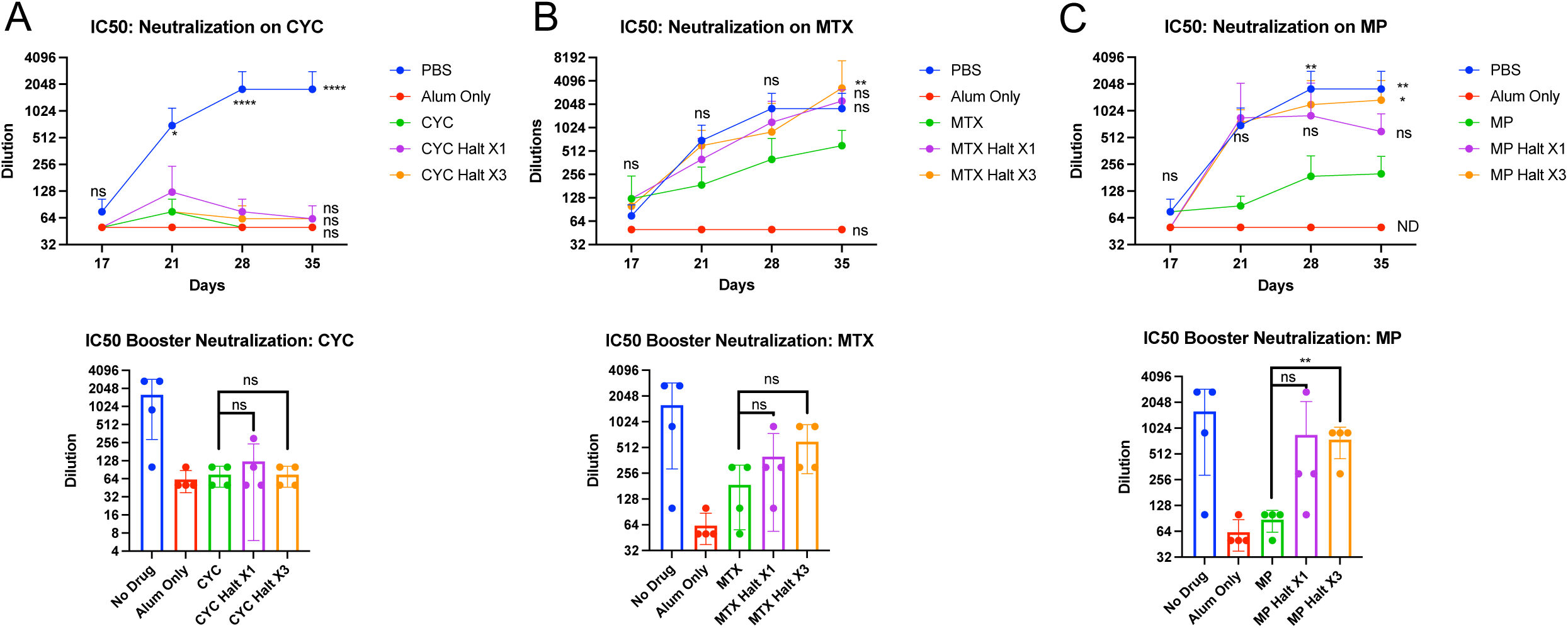
Changes in antibody neutralization efficacy in a VSV-SARS-CoV-2 spike protein system following temporary suspension of immunosuppressant administration in mice. Individual mouse sera samples (n=4) were collected from mouse groups treated or not treated with immunosuppressants at 17, 21, 28, and 35 days, which followed immunization with spike protein (top panels). Virus neutralization was assessed as described in Figure 2. Dilutions needed for 50% neutralization using booster serum (Day 21) samples are indicated at bottom. Noted dilutions are plotted on a Log2 scale. Statistical values were determined as compared to continuous drug treatment (A. CYC; B. MTX; C. MP).

### Immune suppression alters the immune landscape of spike protein-immunized mice

The immunosuppressants used to treat mice have been associated with altering counts and percentages of key immune cell populations in numerous previous research studies(22-30). These changes in immune populations are an important function in human patients that require immune modulation through suppressive therapy. Towards elucidating if this alteration of the immune landscape is directly related to the inhibition of antibody response to spike protein observed in Fig 1B-D and Fig 2, and whether halting treatment at the time of vaccination would affect these changes (including important long-term immune considerations), we euthanized the immunosuppressant-treated (both spike-immunized and control mice) at the experimental endpoint (Day 35) and used flow cytometry to determine the percentages of key immune cell populations within live CD45+ splenocytes and lymphocytes (Fig 5). We observed significant decreases in B cell numbers notably in cyclophosphamide-treated mice. The changes in antibody response to spike protein may therefore be dependent on specific immune cell suppression, particularly of antibody-producing B cells populations. Changes in B cell populations and the consequential skewing of the immune landscape is critical to modulation of the vaccination response to SARS-CoV-2. Temporary suspension of the drugs was sufficient to increase percentages of some cell populations in the CYC group (notably CD4+ T cells) but not for B cells. Strikingly, CYC treatment significantly increased the observed percentage of CD11b+Gr1+ myeloid-derived suppressor cell (MDSC) murine cells within the CD45+ gated populations, indicating another possible mechanism of immune suppression within these mice. Furthermore, within the splenic samples of all CYC-treated mouse groups, we observed a decrease in the percent of TCRD□^+^ cells and an increase in TCRDD□□^+^ immune cells, indicating a skewing of TCR subsets. No significant changes to immune cell populations were observed following temporary suspension of MP or MTX compared to the respective continuous treatment groups, indicating that temporary suspension of these drugs does not alter immune landscape at extended timepoints.

**Figure 5.**
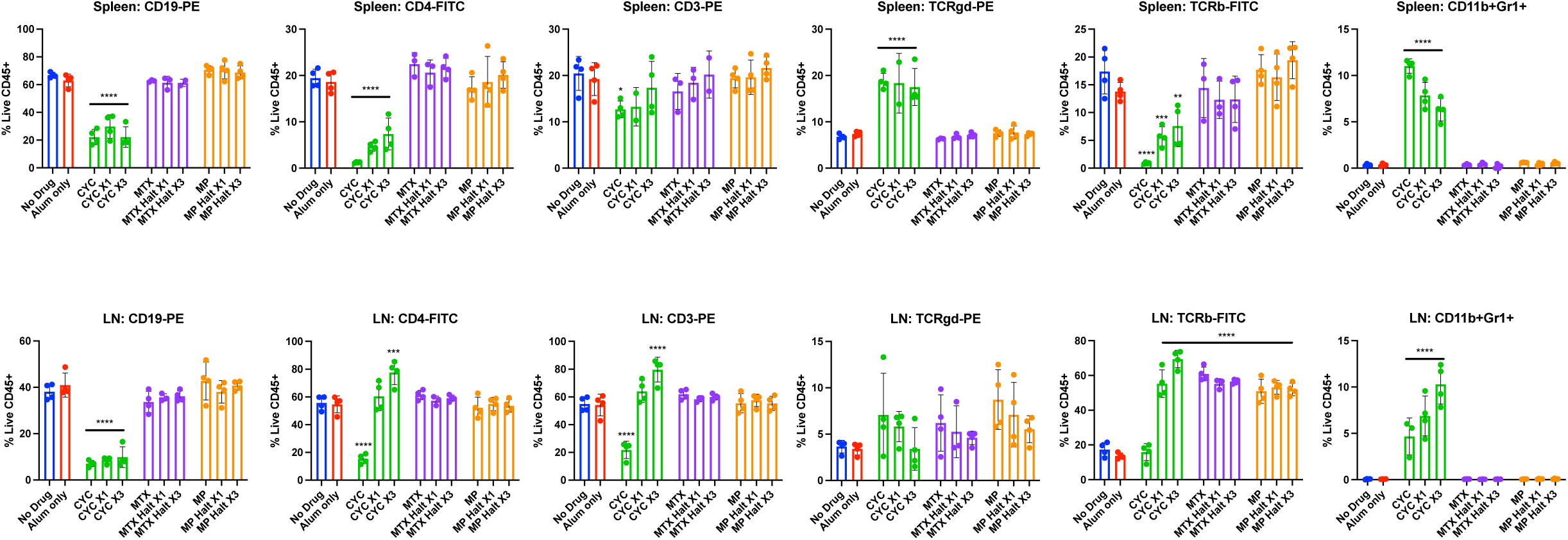
Phenotypic changes in immune cell subpopulations within the spleens and lymph nodes of immunosuppressant treated mice immunized with SARS-CoV-2 spike protein. Immunosuppressant treated, spike protein-immunized mice were euthanized at Day 35, and lymph nodes and spleens were harvested and digested into single-cell suspensions. Tissue samples were stained with indicated conjugated fluorescent antibodies (BioLegend) and analyzed using flow cytometry (Beckman Coulter Cytoflex). Bars represent the mean percentage results obtained from four individual mice (excepting samples in which a minimum number of viable CD45+ cells could not be obtained due to instrument error). Statistical values were determined as compared to No Drug control mice.

## Discussion

The COVID-19 pandemic caused by SARS-COV-2 represented a new paradigm of vaccine and immunotherapy design. Vaccine candidates were designed, tested, and approved for human use in an unprecedented timespan. As of July 2021, the WHO has noted 108 vaccine candidates that have progressed to clinical evaluation. While vaccination against SARS-CoV-2 has resulted in dramatic decreases in COVID-19 case numbers, one growing concern has been the effects of immune suppression on these vaccination efforts. We demonstrate here that commonly used immunosuppressants can significantly impair the antibody response to the spike protein of SARS-CoV-2 in a mouse model. This decrease in antibody titers was observed in all tested immune suppressants, but most notably in cyclophosphamide, a drug that is known to dysregulate immune cell populations and, as our research reinforces, greatly depletes B cell populations. This is particularly noteworthy as previous studies have linked B cell depletion with long-term effects on patient response to vaccination(24) and a relatively high rate of mortality in COVID-19 patients(33). Here, we developed a regimen in which key immunosuppressant treatments were temporarily halted for varied timepoints around the time of vaccinations (Fig 3A). The B cell depletion observed in the cyclophosphamide group was not significantly restored at the time of euthanasia, and antibody levels/functions were not observably altered, suggesting that greater consideration must be given to vaccination and treatment strategies for patients on long-term B cell depletion therapy. We did, however, notice significant increases in the levels of IgG titers against spike protein in the methylprednisolone-treated mouse groups, both when MP was suspended for one timepoint and when suspended for three timepoints. This suggests that, depending on the immunosuppressant and its method of action, an improved response to SARS-CoV-2 vaccination can be achieved with modulation of the drug treatment. However, many additional considerations should be taken into account for future studies and optimization of these treatment regimens, specifically in humans. One important consideration is the ability of SARS-CoV-2 to evade host immune-mediated cell death mechanisms and promoting inflammatory processes. Research has shown the spike protein is highly glycosylated (34, 35); these glycan structures may act as a shield against immune targeting of the SARS-CoV-2 virion. Overcoming this glycan shield to promote immune killing mechanisms (35-37) would thus be an important factor in balancing vaccine strategies with immune modulation.

One limitation of our study is the translation of mouse drug treatment and COVID-19 vaccination into human patients. We selected the drug treatment regimens for our mouse models based on previous research showing drug efficacy with no apparent effects on animal well-being. Also, we used intraperitoneal drug injections for optimal delivery and to minimize variations in individual drug deliveries. This method minimizes potential error in drug delivery, though one potential limit would be faster release into the circulatory system as opposed to subcutaneous or intramuscular injections as sometimes used in humans. Similarly, our generated spike protein was validated in both intraperitoneal and intramuscular injection models, and we used intraperitoneal vaccination in the discussed experiments to minimize the chance for error. While the purified spike protein has not as yet been used as a COVID-19 vaccine in the clinical practice, our validation (Supplementary Fig 2) in addition to the SARS-CoV-2 neutralization experiments (Fig2, Fig 4) show the relevance of this protein to established COVID-19 vaccine practices. Alum was used for spike protein vaccination to optimize immune response and antibody response. However, most currently approved COVID-19 vaccines use different formulations with non-alum adjuvants. While we show the overall importance of regulating immunosuppressant treatment regimens in COVID-19 vaccine strategies, the differences in drug delivery systems, as well as dosages of immunosuppressants, would be an important consideration moving forward in drug optimization.

In their most updated COVID-19 Vaccine Clinical Guidance, American College of Rheumatology recommends timing considerations for immunomodulatory therapy and COVID-19 vaccination (38). Current guidance for MTX and CYC is to withhold their administration for a period during the COVID-19 vaccination. Our findings not only validate this recommendation but also provide experimental evidence for the recommendation. Moreover, our results could serve towards strengthening the current confidence level of task force ranked as “moderate” and guide future clinical studies in humans. Our findings with the other three immunosuppressant drugs would also serve as additional resource to assist with further such clinical guidelines. The COVID-19 pandemic has demonstrated the critical need for constant and progressive research in vaccination and immunotherapeutic strategies. This clarification is especially vital in the context of patients whose health is dependent on immune modulating therapies. We show in this proof-of-principle study that such elucidation is feasible for improving SARS-CoV-2 vaccination response and prognosis for at-risk individuals.

## Materials and Methods

### Production of recombinant SARS-CoV-2 spike protein

The mammalian expression vector, pcDNA3.1+, with the codon-optimized nucleotide sequence of SARS-CoV-2 spike protein was a generous gift from Jarrod Mousa (University of Georgia). The nucleotide sequence of the gene has a “GSAS” substitution at the furin cleavage site (residues 682–685), stabilizing mutations (K986P and V987P), a human rhinovirus 3C protease cleavage site, a T4 foldon trimerization domain and an 8XHisTag. The spike protein was expressed in a serum-free medium by transient transfection of FreeStyle™ 293-F cells and purified by affinity chromatography using Nickel resins. Then, the protein was further purified on a Superdex S200 size exclusion column. The antigenicity and the immunogenicity of the purified recombinant spike protein was validated (Supplementary Fig 1) against verified available recombinant spike protein (BEI Resources, NR52397) and anti-SARS-CoV-2 spike antibody clone A20085C (BioLegend).

### Mice

Eight-week-old female BALB/c mice were obtained from Jackson Laboratories (Bar Harbor, ME) and housed at the University of Georgia. Mice were kept in microisolator cages and handled under biosafety level 2 (BSL2) hoods. For tissue processing and subsequent flow cytometry, mice were euthanized through carbon dioxide in accordance with IACUC guidelines. Serum samples, spleens and lymph nodes were harvested. Cell suspensions were generated through mechanical tissue disruption and collagenase D digestion. Red blood cells were lysed, and samples were filtered through 60 μm nylon filters to obtain single cell suspensions.

All mouse experiments were in compliance with the University of Georgia Institutional Animal Care and Use Committee under an approved animal use protocol. Our animal use protocol adheres to the principles outlined in ***U*.*S. Government Principles for the Utilization and Care of Vertebrate Animals Used in Testing, Research and Training***, the Animal Welfare Act, the ***Guide for the Care and Use of Laboratory Animals***, and the ***AVMA Guidelines for the Euthanasia of Animals***.

### Mouse Treatment and Immunizations

Mouse groups (n=4) were injected with 100μl of vehicle (PBS, or No Drug) or immunosuppressants: cyclophosphamide monohydrate (CYC, Sigma Aldrich C0768, 60mg/kg every other day), leflunomide (LEF, Sigma Aldrich L5025, 20mg/kg every other day), methotrexate (MTX, Sigma Aldrich M9929, 1mg/kg every other day), 6a-methylprednisolone (MP, Sigma Aldrich M0639, 20mg/kg every other day), and mycophenolate mofetil (MM, Tocris 4102, 40mg/kg every other day). These dosages were selected based on previous established murine research models for each of the respective drug treatments (22-30) to ensure efficacy. Treatments were started at seven days before immunization and administered intraperitoneally (IP). Mice were immunized with SARS-CoV-2 spike protein at 0.5 μg/50μl alum (InvivoGen, stock concentration aluminum 10 μg/μl)/mouse intraperitoneally on days 0 and 14 (Fig 1A). Alum only mouse groups received 50μl alum alone intraperitoneally. For experiments in which immunosuppressant regimens were temporarily halted at the time of vaccinations, mouse groups received continuous treatments starting at day -8 except at the day of immunization (Halt X1) or except for the two days prior to, two days following, and day of immunization (Halt X3) (Fig 3A).

### ELISA

Mice were bled from the tail vein 14 days after initial immunization of spike protein or alum only, and days 17, 21, 28, and 35 (post boost). Sera samples were stored at -20oC until time of ELISA (after endpoint). Spike protein-specific antibodies in serum were detected by ELISA in 384-well plates coated with 0.5 μg/ml of spike protein for 24 hours. All mouse sera samples (different drug groups and different sera collection dates) were tested on the same 384-well plate to eliminate plate-to-plate variation. One 384-well plate was used to test one dilution and one antibody (IgG or IgM). Four immune sera per group were used in all ELISA experiments. Anti-IgG-AP (Southern BioTech 1030-04) and anti-IgM-AP (Southern BioTech 1020-04) were used to detect antibodies. Absorbance was measured at 405nm. Regression curves based on dilutions were used to determine IgG and IgM titer levels at an absorbance of 0.5. Shown are calculated mouse titers (reciprocal dilutions) at absorbance of 0.5 for individual mice and average values with statistical significance indicated. For ELISA data in Fig. 1B and Fig 3B-D (upper panels), input was OD values of individual sera (n=4) diluted 3200 times at 405nm absorbance and significance.

### Flow Cytometry

Immunosuppressant treated (including spike-immunized and control) mice were euthanized at Day 35, and spleens and lymph nodes were harvested and digested into single-cell suspensions. Cells were stained in PBS with TruStain fcX (BioLegend, Cat. No. 101320) to reduce non-specific antibody binding. Cells were then stained with the following antibodies and stains (in multiple sets to prevent fluorophore overlap): CD4-FITC (BioLegend), CD8-PECy5 (BioLegend), CD3-PE(BioLegend), CD45-APCCy7 (BioLegend), CD19-PE (BioLegend), Ghost Red 710 (Tonbo 13-0871-T100), TCR□-FITC (BioLegend), TCR□□-PE (BioLegend), Gr1-PECy5 (BioLegend), and CD11b-FITC (BioLegend). Samples were washed and analyzed with flow cytometry (Beckman Coulter CytoFLEX). Isotype control antibody-stained samples were used as negative staining controls where appropriate. Flow cytometry data was analyzed using FlowJo Single Cell Analysis Software (Treestar, Inc., Ashland, Oregon) with gating strategies shown in Supplementary Figure 2. Briefly, live CD45+ immune cells were gated and percentages of immune populations were derived from these gates.

### Neutralizing antibody assay

Individual mouse sera (n=4) were heat inactivated (56°C, 30 min) and serial diluted in Dulbecco’s Modified Eagle Medium (DMEM). Diluted sera were mixed with approximated 400 infectious particles of a recombinant vesicular stomatitis virus encoding the SARS-CoV-2 spike protein and nano-luciferase (room temperature, 30 min)(31, 32). To improve spike incorporation onto rVSV particles the cytoplasmic tail was removed. The spike encodes for the Wuhan isolate with the D614G amino acid change (rVSVΔG/SARS-CoV-2-SΔ21-D614G-NLucP). The virus-sera mixture was added to three replicate wells of Vero cells in a 96-well plate and incubated for 24 hr at 37°C. Virus infection was monitored by luciferase activity. Cells were lysed with Nano-Glo Luciferase Assay system (Promega), lysates were transferred to white plates and luminescence was read in a GloMax® Discover Microplate Reader (Promega). Neutralization activity was determined by comparing the signal from the wells infected with virus lacking antibody to the different sera dilutions. The antibody dilution that reduced luciferase signal by 50% were reported as IC50 concentrations. Dilutions were indicated on a log-2 scale.

### Statistical Analysis

GraphPad Prism v8 was used for statistical analyses. For ELISA titer data, flow cytometry immunophenotyping, and single timepoint neutralization, ordinary one-way ANOVA with Dunnett’s multiple comparisons tests was used to determine statistical significance between experimental groups in each of the applicable experimental models. For multiple timepoint ELISA and neutralization experiments, 2way ANOVA with Tukey’s multiple comparisons tests was used to determine statistical significance between experimental groups in each of the applicable experimental models. For direct comparison between continuous drug treatment and Halt X1 or Halt X3 in Figure 4, unpaired parametric two-tailed t test was used. Statistical comparisons were made with “No Drug” as the control group for Figures 1 and 2. Comparisons were made between continuous treatments with the respective drugs (CYC, MTX, and MP) as the control groups in Figures 3 and 4. Significance is indicated on each graph based on p value: >0.05 = ns; <.05 = *; <0.01 = **; <0.001 = ***; <0.0001 = ****. 95% CI ranges and p values are listed in Supplementary Table 3. Furthermore, effect sizes were calculated with No Drug (Fig1-2) or continuous drug treatment (Fig3-4) using Excel to find Cohen’s d (standardized mean difference). The effect size threshold was determined to be small (d<0.5), medium (d<0.8), large (d<1.30) or very large(d>1.30). These values have been included in Supplementary Tables 3-6.

## Acknowledgments

This work was supported by National Institutes of Health grants R01AI123383 (FYA), R01AI152766 (FYA).

