## Supplemental Figures and Tables for "Modulation of immunosuppressant drug treatment to improve SARS-CoV-2 vaccine efficacy in mice"

### Supplementary Figures and Tables

**Supplementary Figure 1.**

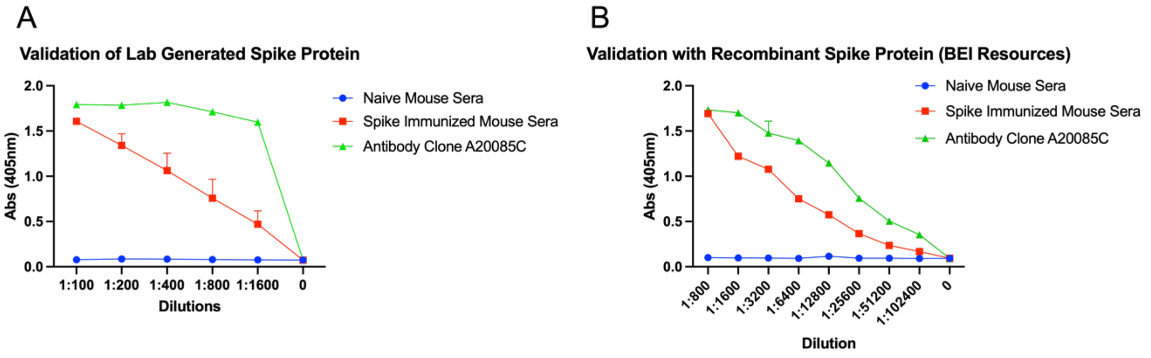

**Supplementary Figure 1. Validation of recombinantly generated SARS-CoV-2 spike protein and mouse immunizations.** Mice were immunized with 0.5 μg spike protein generated in our lab as described in the Methods. Mice were boosted with spike protein 14 days after initial immunization, and serum samples were obtained 14 days after booster immunization. A. To validate the antigenicity of our recombinant spike protein, ELISA plates were coated with our recombinant spike protein overnight. ELISA was performed using naïve or spike-immunized mouse serum, in tandem with commercially available and validated anti-SARS-CoV-2 spike antibody (BioLegend, clone A20085C starting dilution of 0.5μg/100μl/well). Mouse IgG-AP was used as secondary and absorbances were detected at 405nm. B. To validate the immunogenicity of our recombinant spike protein in comparison with validated, available spike protein, we performed an ELISA with commercially available recombinant spike protein (BEI Resources NR52397) coated ELISA plates. Sera from recombinant spike (our lab)-immunized mice and naïve mice were compared to commercially available and validated anti-SARS-CoV-2 spike antibody (BioLegend, clone A20085C, starting dilution of 0.5μg/100μl/well). Mouse IgG-AP was used as secondary and absorbances were detected at 405nm.

**Supplementary Figure 2.**

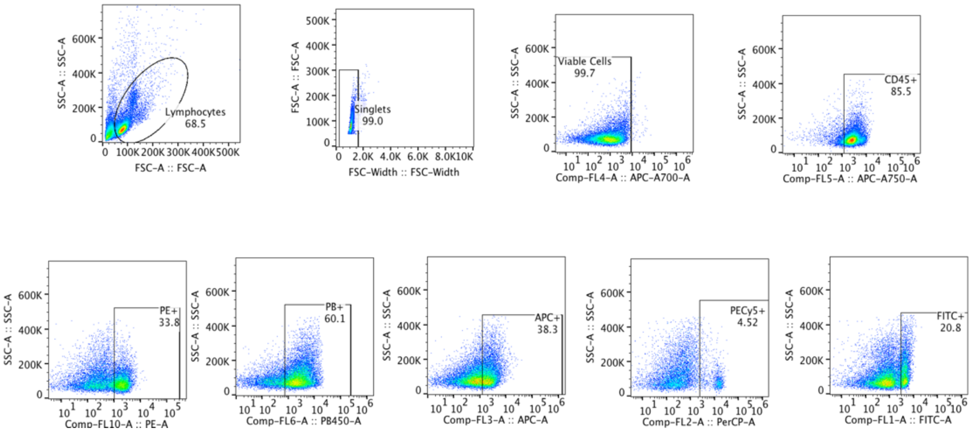

**Supplementary Figure 2. Supplementary Figure 2: Flow cytometry gating strategy.** Lymph nodes and spleens were harvested and digested into single-cell suspensions. Tissue samples were stained with indicated conjugated fluorescent antibodies (BioLegend) and analyzed using flow cytometry (Beckman Coulter Cytoflex). Single cells were gated based on FSC and SSC as indicated. Gates were assigned based on single staining and negative isotype control-stained populations.

**Supplementary Table 1.**

| Treatment | Regimen | Immunize | Number Mice | Dosage |
|---|---|---|---|---|
| No Drug/PBS | Continuous | Spike protein + alum | 5 | |
| Methylprednisolone | Continuous | Spike protein + alum | 4 | 20mg/kg Every other day |
| Methotrexate | Continuous | Spike protein + alum | 4 | 1mg/kg Every other day |
| Cyclophosphamide | Continuous | Spike protein + alum | 4 | 60mg/kg Every other day |
| Leflunomide | Continuous | Spike protein + alum | 4 | 20 mg/kg Every other day |
| Mycophenolate mofetil | Continuous | Spike protein + alum | 4 | 40 mg/kg Every other day |
| Hydroxychloroquine | Continuous | Spike protein + alum | 4 | 40 mg/kg Every day |

**Supplementary Table 1. Supplementary Table 1: Immunosuppressant administration.** Each drug was administered intraperitoneally at 100μl/mouse at the indicated times starting at 7 days before initial spike protein immunization and continuously up through time of euthanasia.

**Supplementary Table 2.**

| Treatment | Regimen | Immunize | Number Mice | Dosage |
|---|---|---|---|---|
| No Drug/PBS | Continuous | 1. Spike protein + alum<br>2. Alum only | 4<br>4 | |
| Methylprednisolone | 1. Continuous<br>2. Halt on Days -2, 0, +2<br>3. Halt on Day 0 | Spike protein + alum | 4<br>4<br>4 | 20mg/kg<br>Every other day |
| Methotrexate | 1. Continuous<br>2. Halt on Days -2, 0, +2<br>3. Halt on Day 0 | Spike protein + alum | 4<br>4<br>4 | 1mg/kg<br>Every other day |
| Cyclophosphamide | 1. Continuous<br>2. Halt on Days -2, 0, +2<br>3. Halt on Day 0 | Spike protein + alum | 4<br>4<br>4 | 60mg/kg<br>Every other day |

**Supplementary Table 2. Supplementary Table 2: Immunosuppressant administration with temporary suspension of treatment.** Each drug was administered intraperitoneally at 100μl/mouse at the indicated times starting at 8 days before initial spike protein immunization and continuously up through time of euthanasia, except at indicated timepoints.

**Supplementary Table 3.**

| Comparison | Mean Diff. | 95% CI of diff. | Summary | Adjusted P Value |
|---|---|---|---|---|
| Figure 1B Time Course | | | | |
| Day 21 | | | | |
| No Drug vs. Alum only | 0.4408 | 0.3977 to 0.4839 | **** | <0.0001 |
| No Drug vs. CYC | 0.4413 | 0.4028 to 0.4799 | **** | <0.0001 |
| No Drug vs. MTX | 0.3643 | 0.3258 to 0.4029 | **** | <0.0001 |
| No Drug vs. MP | 0.278 | 0.2395 to 0.3165 | **** | <0.0001 |
| No Drug vs. MM | 0.06567 | 0.02712 to 0.1042 | *** | 0.0002 |
| No Drug vs. LEF | 0.1743 | 0.1358 to 0.2129 | **** | <0.0001 |
| Day 28 | | | | |
| No Drug vs. Alum only | 0.5762 | 0.5331 to 0.6193 | **** | <0.0001 |
| No Drug vs. CYC | 0.5817 | 0.5431 to 0.6202 | **** | <0.0001 |
| No Drug vs. MTX | 0.4833 | 0.4448 to 0.5219 | **** | <0.0001 |
| No Drug vs. MP | 0.3473 | 0.3088 to 0.3859 | **** | <0.0001 |
| No Drug vs. MM | 0.1333 | 0.09478 to 0.1719 | **** | <0.0001 |
| No Drug vs. LEF | 0.3053 | 0.2668 to 0.3439 | **** | <0.0001 |
| Day 35 | | | | |
| No Drug vs. Alum only | 0.5353 | 0.4922 to 0.5784 | **** | <0.0001 |
| No Drug vs. CYC | 0.5383 | 0.4998 to 0.5769 | **** | <0.0001 |
| No Drug vs. MTX | 0.368 | 0.3295 to 0.4065 | **** | <0.0001 |
| No Drug vs. MP | 0.3273 | 0.2888 to 0.3659 | **** | <0.0001 |
| No Drug vs. MM | 0.1467 | 0.1081 to 0.1852 | **** | <0.0001 |
| No Drug vs. LEF | 0.245 | 0.2065 to 0.2835 | **** | <0.0001 |

| Comparison | Mean Diff. | 95% CI of diff. | Summary | Adjusted P Value | Effect Size (Cohen's d) | Effect Size Threshold |
|---|---|---|---|---|---|---|
| Figure 1C Booster | | | | | | |
| PBS vs. Alum only | 10414 | 7118 to 13710 | **** | <0.0001 | -4.597339412 | Very Large |
| PBS vs. CYC | 10489 | 7193 to 13785 | **** | <0.0001 | -4.629069676 | Very Large |
| PBS vs. MTX | 9839 | 6543 to 13135 | **** | <0.0001 | -4.342295079 | Very Large |
| PBS vs. MP | 9241 | 5945 to 12537 | **** | <0.0001 | -3.616011959 | Very Large |
| PBS vs. MM | 6656 | 3360 to 9953 | **** | <0.0001 | -2.704941817 | Very Large |
| PBS vs. LEF | 8363 | 5066 to 11659 | **** | <0.0001 | -3.120982747 | Very Large |
| Figure 1C Endpoint | | | | | | |
| PBS vs. Alum only | 35857 | 20825 to 50890 | **** | <0.0001 | -3.264025328 | Very Large |
| PBS vs. CYC | 36087 | 21055 to 51120 | **** | <0.0001 | -3.28610168 | Very Large |
| PBS vs. MTX | 30560 | 15528 to 45593 | **** | <0.0001 | -2.782808562 | Very Large |
| PBS vs. MP | 30389 | 15356 to 45421 | **** | <0.0001 | -2.481970605 | Very Large |
| PBS vs. MM | 23980 | 8948 to 39013 | ** | 0.0012 | -1.989205938 | Very Large |
| PBS vs. LEF | 29088 | 14055 to 44120 | *** | 0.0001 | -2.41221589 | Very Large |
| Figure 1D Booster | | | | | | |
| No Drug vs. Alum only | 3064 | 1684 to 4445 | **** | <0.0001 | -3.772390939 | Very Large |
| No Drug vs. CYC | 2658 | 1278 to 4039 | *** | 0.0002 | -3.278275916 | Very Large |
| No Drug vs. MTX | 163.6 | -1217 to 1544 | ns | 0.9983 | -0.201709004 | Small |
| No Drug vs. MP | 1613 | 232.6 to 2993 | * | 0.0181 | -1.876429265 | Very Large |
| No Drug vs. MM | -536.4 | -1917 to 843.7 | ns | 0.7667 | 0.568792029 | Medium |
| No Drug vs. LEF | 874.9 | -505.3 to 2255 | ns | 0.3313 | -0.816853605 | Large |
| Figure 1D Endpoint | | | | | | |
| PBS vs. Alum only | 35857 | 20825 to 50890 | **** | <0.0001 | -2.514073525 | Very Large |
| PBS vs. CYC | 36087 | 21055 to 51120 | **** | <0.0001 | -2.156649396 | Very Large |
| PBS vs. MTX | 30560 | 15528 to 45593 | **** | <0.0001 | -1.232620086 | Large |
| PBS vs. MP | 30389 | 15356 to 45421 | **** | <0.0001 | -0.949748227 | Large |
| PBS vs. MM | 23980 | 8948 to 39013 | ** | 0.0012 | -0.246501342 | Small |
| PBS vs. LEF | 29088 | 14055 to 44120 | *** | 0.0001 | -0.220681398 | Small |

**Supplementary Table 3. Statistical evaluation of changes in antibody levels to SARS-CoV-2 spike protein following immunosuppressant administration.** ANOVA multiple comparisons tests were performed on the sample groups as described in Figure1. Shown are mean of differences, 95% CI of differences, adjusted P values, and summary of statistical significance based on P values. For titer calculations, effect size calculations were performed to obtain a standardized mean difference (Cohen's d) as well as its interpreted effect size threshold.

**Supplementary Table 4.**

| Comparison | Mean Diff. | 95% CI of diff. | Summary | Adjusted P Value | Effect Size (Cohen's d) | Effect Size Threshold |
|---|---|---|---|---|---|---|
| Figure 2B | | | | | | |
| No Drug vs. Alum only | 4900 | 1506 to 8294 | ** | 0.0034 | -1.867429038 | Very Large |
| No Drug vs. CYC | 4900 | 1506 to 8294 | ** | 0.0034 | -1.867429038 | Very Large |
| No Drug vs. MTX | 4867 | 1201 to 8533 | ** | 0.007 | -1.854725439 | Very Large |
| No Drug vs. MP | 4100 | 705.9 to 7494 | * | 0.0146 | -1.482358128 | Very Large |
| No Drug vs. MM | 4650 | 493.1 to 8807 | * | 0.025 | -1.772152046 | Very Large |
| No Drug vs. LEF | 3363 | -31.63 to 6757 | ns | 0.0528 | -1.206265899 | Large |

**Supplementary Table 4. Statistical evaluation of changes in antibody neutralization efficacy in a VSV-SARS-CoV-2 spike protein system.** ANOVA multiple comparisons tests were performed on the sample groups as described in Figure2. Shown are mean of differences, 95% CI of differences, adjusted P values, and summary of statistical significance based on P values. For neutralization dilution values at Day 21, effect size calculations were performed to obtain a standardized mean difference (Cohen's d) as well as its interpreted effect size threshold, comparing each immunosuppressant drug group (as well as Alum only) to the No Drug control group.

**Supplementary Table 5.**

| Comparison | Mean Diff. | 95% CI of diff. | Summary | Adjusted P Value |
|---|---|---|---|---|
| Figure 3B | | | | |
| Day 21 | | | | |
| CYC vs. No Drug | -0.7189 | -0.7726 to -0.6651 | **** | <0.0001 |
| CYC vs. Alum only | -0.0065 | -0.06024 to 0.04724 | ns | 0.9944 |
| CYC vs. CYC Halt X1 | -0.0115 | -0.06524 to 0.04224 | ns | 0.9548 |
| CYC vs. CYC Halt X3 | -0.005 | -0.05874 to 0.04874 | ns | 0.9978 |
| Day 28 | | | | |
| CYC vs. No Drug | -0.732 | -0.7857 to -0.6783 | **** | <0.0001 |
| CYC vs. Alum only | -0.002125 | -0.05587 to 0.05162 | ns | 0.9999 |
| CYC vs. CYC Halt X1 | -0.014 | -0.06774 to 0.03974 | ns | 0.9135 |
| CYC vs. CYC Halt X3 | 0.003125 | -0.05062 to 0.05687 | ns | 0.9997 |
| Day 35 | | | | |
| CYC vs. No Drug | -0.7333 | -0.7870 to -0.6795 | **** | <0.0001 |
| CYC vs. Alum only | -0.000625 | -0.05437 to 0.05312 | ns | >0.9999 |
| CYC vs. CYC Halt X1 | 0.000625 | -0.05312 to 0.05437 | ns | >0.9999 |
| CYC vs. CYC Halt X3 | 0.01087 | -0.04287 to 0.06462 | ns | 0.9627 |
| Figure 3C | | | | |
| Day 21 | | | | |
| MTX vs. No Drug | -0.4423 | -0.5833 to -0.3012 | **** | <0.0001 |
| MTX vs. Alum only | 0.2701 | 0.1291 to 0.4111 | **** | <0.0001 |
| MTX vs. MTX Halt X1 | -0.1444 | -0.2854 to -0.003353 | * | 0.0432 |
| MTX vs. MTX Halt X3 | -0.2155 | -0.3565 to -0.07448 | ** | 0.0011 |
| Day 28 | | | | |
| MTX vs. No Drug | -0.3729 | -0.5139 to -0.2319 | **** | <0.0001 |
| MTX vs. Alum only | 0.357 | 0.2160 to 0.4980 | **** | <0.0001 |
| MTX vs. MTX Halt X1 | -0.1564 | -0.2974 to -0.01535 | * | 0.0251 |
| MTX vs. MTX Halt X3 | -0.189 | -0.3300 to -0.04798 | ** | 0.0049 |
| Day 35 | | | | |
| MTX vs. No Drug | -0.278 | -0.4190 to -0.1370 | **** | <0.0001 |
| MTX vs. Alum only | 0.4546 | 0.3136 to 0.5956 | **** | <0.0001 |
| MTX vs. MTX Halt X1 | -0.06662 | -0.2076 to 0.07440 | ns | 0.5837 |
| MTX vs. MTX Halt X3 | -0.09012 | -0.2311 to 0.05090 | ns | 0.3201 |
| Figure 3D | | | | |
| MP vs. No Drug | -0.5183 | -0.6510 to -0.3855 | **** | <0.0001 |
| MP vs. Alum only | 0.1941 | 0.06138 to 0.3269 | ** | 0.0019 |
| MP vs. MP Halt X1 | -0.34 | -0.4727 to -0.2073 | **** | <0.0001 |
| MP vs. MP Halt X3 | -0.36 | -0.4927 to -0.2273 | **** | <0.0001 |
| Day 28 | | | | |
| MP vs. No Drug | -0.4229 | -0.5556 to -0.2901 | **** | <0.0001 |
| MP vs. Alum only | 0.307 | 0.1743 to 0.4397 | **** | <0.0001 |
| MP vs. MP Halt X1 | -0.1923 | -0.3250 to -0.05951 | ** | 0.0021 |
| MP vs. MP Halt X3 | -0.2868 | -0.4195 to -0.1540 | **** | <0.0001 |
| Day 35 | | | | |
| MP vs. No Drug | -0.3895 | -0.5222 to -0.2568 | **** | <0.0001 |
| MP vs. Alum only | 0.3431 | 0.2104 to 0.4759 | **** | <0.0001 |
| MP vs. MP Halt X1 | -0.2155 | -0.3482 to -0.08276 | *** | 0.0005 |
| MP vs. MP Halt X3 | -0.2308 | -0.3635 to -0.09801 | *** | 0.0002 |

| Comparison | Mean Diff. | 95% CI of diff. | Summary | Adjusted P Value | Effect Size (Cohen's d) | Effect Size Threshold |
|---|---|---|---|---|---|---|
| Figure 3B Titers | | | | | | |
| CYC vs. No Drug | -10905 | -11877 to -9933 | **** | <0.0001 | | |
| CYC vs. Alum only | 650.7 | -321.5 to 1623 | ns | 0.2468 | | |
| CYC vs. CYC Halt X1 | -410.3 | -1383 to 561.9 | ns | 0.6162 | -0.925387596 | Large |
| CYC vs. CYC Halt X3 | -448.4 | -1421 to 523.8 | ns | 0.5461 | -0.845383449 | Large |
| Figure 3C Titers | | | | | | |
| MTX vs. No Drug | -9623 | -11554 to -7692 | **** | <0.0001 | | |
| MTX vs. Alum only | 1933 | 2.130 to 3863 | * | 0.0497 | | |
| MTX vs. MTX Halt X1 | -2664 | -4594 to -733.1 | ** | 0.0065 | -1.838339371 | Very Large |
| MTX vs. MTX Halt X3 | -2816 | -4747 to -885.9 | ** | 0.0042 | -8.445216204 | Very Large |
| Figure 3D Titers | | | | | | |
| MP vs. No Drug | -10084 | -12735 to -7432 | **** | <0.0001 | | |
| MP vs. Alum only | 1472 | -1179 to 4123 | ns | 0.3924 | | |
| MP vs. MP Halt X1 | -3729 | -6380 to -1078 | ** | 0.0056 | -2.294945283 | Very Large |
| MP vs. MP Halt X3 | -4870 | -7521 to -2219 | *** | 0.0006 | -3.448740902 | Very Large |

**Supplementary Table 5. Statistical evaluation of changes in antibody titers to SARS-CoV-2 spike protein following continuous or temporarily suspended immunosuppressant administration.** ANOVA multiple comparisons tests were performed on the sample groups as described in Figure3. Shown are mean of differences, 95% CI of differences, adjusted P values, and summary of statistical significance based on P values. For titer calculations, effect size calculations were performed to obtain a standardized mean difference (Cohen's d) as well as its interpreted effect size threshold, comparing Halt X1 and X3 treatments to the continuous drug group.

**Supplementary Table 6.**

| Comparison | Mean Diff. | 95% CI of diff. | Summary | Adjusted P Value | Effect Size (Cohen's d) | Effect Size Threshold |
|---|---|---|---|---|---|---|
| Figure 4A Time Course | | | | | | |
| 21 | | | | | | |
| CYC vs. PBS | -625 | -1232 to -18.34 | * | 0.0415 | | |
| CYC vs. Alum Only | 25 | -581.7 to 631.7 | ns | 0.9999 | | |
| CYC vs. CYC Halt X1 | -50 | -656.7 to 556.7 | ns | 0.9987 | | |
| CYC vs. CYC Halt X3 | 0 | -606.7 to 606.7 | ns | >0.9999 | | |
| 28 | | | | | | |
| CYC vs. PBS | -1750 | -2357 to -1143 | **** | <0.0001 | | |
| CYC vs. Alum Only | 0 | -606.7 to 606.7 | ns | >0.9999 | | |
| CYC vs. CYC Halt X1 | -25 | -631.7 to 581.7 | ns | 0.9999 | | |
| CYC vs. CYC Halt X3 | -12.5 | -619.2 to 594.2 | ns | >0.9999 | | |
| 35 | | | | | | |
| CYC vs. PBS | -1750 | -2357 to -1143 | **** | <0.0001 | | |
| CYC vs. Alum Only | 0 | -606.7 to 606.7 | ns | >0.9999 | | |
| CYC vs. CYC Halt X1 | -12.5 | -619.2 to 594.2 | ns | >0.9999 | | |
| CYC vs. CYC Halt X3 | -12.5 | -619.2 to 594.2 | ns | >0.9999 | | |
| Figure 4A Dilutions | | | | | | |
| CYC vs. CYC Halt X1 | 50.00 ± 61.24 | -99.84 to 199.8 | ns | 0.4454 | -0.577350269 | Medium |
| CYC vs. CYC Halt X3 | 0.000 ± 20.41 | -49.95 to 49.95 | ns | >0.9999 | 0 | Small |
| Figure 4B TimeCourse | | | | | | |
| 21 | | | | | | |
| MTX vs. PBS | -512.5 | -2196 to 1171 | ns | 0.8596 | | |
| MTX vs. Alum Only | 137.5 | -1546 to 1821 | ns | 0.9987 | | |
| MTX vs. MTX Halt X1 | -212.5 | -1896 to 1471 | ns | 0.9933 | | |
| MTX vs. MTX Halt X3 | -412.5 | -2096 to 1271 | ns | 0.9283 | | |
| 28 | | | | | | |
| MTX vs. PBS | -1400 | -3083 to 283.4 | ns | 0.1291 | | |
| MTX vs. Alum Only | 350 | -1333 to 2033 | ns | 0.9586 | | |
| MTX vs. MTX Halt X1 | -800 | -2483 to 883.4 | ns | 0.577 | | |
| MTX vs. MTX Halt X3 | -500 | -2183 to 1183 | ns | 0.8694 | | |
| 35 | | | | | | |
| MTX vs. PBS | -1200 | -2886 to 486.4 | ns | 0.2339 | | |
| MTX vs. Alum Only | 550 | -1136 to 2236 | ns | 0.8322 | | |
| MTX vs. MTX Halt X1 | -1650 | -3336 to 36.37 | ns | 0.0569 | | |
| MTX vs. MTX Halt X3 | -2700 | -4521 to -878.5 | ** | 0.0017 | | |
| Figure 4B Dilutions | | | | | | |
| MTX vs. MTX Halt X1 | 212.5 ± 185.3 | -240.8 to 665.8 | ns | 0.295 | -0.811057872 | Large |
| MTX vs. MTX Halt X3 | 412.5 ± 185.3 | -40.83 to 865.8 | ns | 0.0676 | -1.574406457 | Very Large |

| Comparison | Mean Diff. | 95% CI of diff. | Summary | Adjusted P Value | Effect Size (Cohen's d) | Effect Size Threshold |
|---|---|---|---|---|---|---|
| Figure 4C TimeCourse | | | | | | |
| 21 | | | | | | |
| MP vs. PBS | -612.5 | -1693 to 468.1 | ns | 0.4238 | | |
| MP vs. Alum Only | 37.5 | -1043 to 1118 | ns | 0.9999 | | |
| MP vs. MP Halt X1 | -762.5 | -1843 to 318.1 | ns | 0.2391 | | |
| MP vs. MP Halt X3 | -662.5 | -1743 to 418.1 | ns | 0.3548 | | |
| 28 | | | | | | |
| MP vs. PBS | -1613 | -2693 to -531.9 | ** | 0.0016 | | |
| MP vs. Alum Only | 137.5 | -943.1 to 1218 | ns | 0.9932 | | |
| MP vs. MP Halt X1 | -712.5 | -1793 to 368.1 | ns | 0.293 | | |
| MP vs. MP Halt X3 | -1013 | -2093 to 68.15 | ns | 0.0725 | | |
| 35 | | | | | | |
| MP vs. PBS | -1600 | -2681 to -519.4 | ** | 0.0017 | | |
| MP vs. Alum Only | 150 | -930.6 to 1231 | ns | 0.9903 | | |
| MP vs. MP Halt X1 | -400 | -1481 to 680.6 | ns | 0.7603 | | |
| MP vs. MP Halt X3 | -1150 | -2231 to -69.35 | * | 0.0336 | | |
| Figure 4C Dilutions | | | | | | |
| MP vs. MP Halt X1 | 762.5 ± 618.6 | -751.1 to 2276 | ns | 0.2638 | -0.871606468 | Large |
| MP vs. MP Halt X3 | 662.5 ± 150.5 | 294.2 to 1031 | ** | 0.0046 | -3.112267163 | Very Large |

**Supplementary Table 6. Statistical evaluation of changes in antibody neutralization efficacy in a VSV-SARS-CoV-2 spike protein system following temporary suspension of immunosuppressant administration in mice.** ANOVA multiple comparisons and unpaired t tests were performed on the sample groups as described in Figure4. Shown are mean of differences, 95% CI of differences, adjusted P values, and summary of statistical significance based on P values. For neutralization dilution values at Day 21, effect size calculations were performed comparing Halt X1 and X3 treatments to the continuous drug group to obtain a standardized mean difference (Cohen's d) as well as its interpreted effect size threshold.
